# Neurodegeneration-related beta-amyloid as autocatabolism-attenuator in a *micro-in vivo* system

**DOI:** 10.1101/2020.04.27.063115

**Authors:** Evelin Balazs, Zita Galik-Olah, Bence Galik, Zsolt Bozso, Janos Kalman, Zsolt Datki

## Abstract

Investigation of human neurodegeneration-related aggregates of beta-amyloid 1-42 (Aβ42) on bdelloid rotifers is a novel interdisciplinary approach in life sciences. We reapplied an organ size-based *in vivo* monitoring system, exploring the autocatabolism-related alterations evoked by Aβ42, in a glucose-supplemented starvation model. The experientially easy-to-follow size reduction of the bilateral reproductive organ (germovitellaria) in fasted rotifers was rescued by Aβ42, serving as a nutrient source- and peptide sequence-specific attenuator of the organ shrinkage phase and enhancer of the regenerative one including egg reproduction. Recovery of the germovitellaria was significant in comparison with the greatly shrunken form. In contrast to the well-known neurotoxic Aβ42 (except the bdelloids) with specific regulatory roles, the artificially designed scrambled version (random order of amino acids) was inefficient in autocatabolism attenuation, behaving as negative control. This native Aβ42-related modulation of the ‘functionally reversible organ shrinkage’ can be a potential experiential and supramolecular marker of autocatabolism *in vivo*.

## Introduction

Rotifers are widely accepted animal models of aging-, metabolism-, starvation-, pharmacology- and *micro-in vivo* OMICS research and methodologies (Macsai et al., 2019; Snell, 2014). Bdelloids, such as *Philodina* or *Adineta* species have high tolerance to normal environmental changes due to their capability of adaptive phenotypic plasticity (van Cleave, 1932; Azevedo and Leroi, 2001). Marotta et al. (2012) found that the organs of these animals appeared to be compressed during starvation. Although the germovitellaria (the combined site of the germinarium and vitellarium glands) significantly showed reduction in size and in fine structure under caloric restriction. These organ tissues could also function as nutrient storage; therefore, their content is used by rotifers via autophagy (Mizushima and Komatsu, 2011). In acidic vesicular organelles (AVOs) detection, acidotropic dyes are applied such as acridine orange (AO) or neutral red (NR). These indicators show good correlation with each other (Morishita et al., 2017) and in some cases these fluorescent probes, without cross-linking, are more promising quantitative approaches than immunofluorescence to evaluate the late phase of autocatabolism. The vacuolar-type H^+^-ATPase inhibitors (e.g. Concanamycin A) can hinder the catabolic processes of metabolic autophagy (Goto-Yamada et al., 2019). The direct connection between anatomical changes (e.g. organ shrinkage) and autophagy has been proved in rotifers (Marotta et al., 2012; Cervellione et al., 2017). These phylogenetically ‘simple’ animals are inappropriate to model higher, species-specific (e.g. human) physiological processes (e.g. neurodegeneration); nevertheless, they are suitable for demonstrating various interdisciplinary concepts (Ricci and Boschetti, 2003; Ramulu et al., 2012). An adequate example of this is that the bdelloids are able to exceedingly catabolize the highly resistant peptide- or protein aggregates (well-known neurotoxins), such as beta-amyloid (Aβ), alpha-synuclein and scrapie prions under extreme conditions (e.g. starvation). Investigation of neurodegeneration-related Aβ42 on rotifers is the central research field of our team. In an exceptional way, the human-type aggregates are potential nutrient sources to bdelloid rotifers (Datki et al., 2018); however, the general foods of these animals are aggregated organic masses in their natural habitat (Fontaneto et al., 2011). By the phylogenetically selected ability of bdelloids they are able to use conglomerates and aggregates as an energy source. The rotifer-aggregates-connected research is rather a new topic, there is no relevant data about the Aβ-induced signalizations and regulations in these *micro-in vivo* entities. In this present study, our aims were to reveal the potential regulatory effects of aggregated Aβ42 in relation to the autocatabolism in rotifers.

## Materials and methods

### The invertebrate models

The experiments were performed on *Philodina acuticornis* and *Adineta vaga* species; therefore, no specific ethical permission was needed according to the current international regulations. They were obtained from a Hungarian aquavaristique with origins from an agricultural farm in Szarvas, Hungary. The species have been maintained in standard laboratory conditions for 6 years. (Olah et al., 2017)

### Materials

The Aβ42 and its scrambled isoform (S-Aβ42: LKAFDIGVEYNKVGEGFAISHGVAHLDVSMFGEIGRVDVHQA) were prepared at the Department of Medical Chemistry, University of Szeged, Hungary. The concentrations of the stock solutions were 1 mg/mL in distilled water; the aggregation time was 3h (25 °C, pH 3.5); the neutralization (to pH 7.5) was performed with NaOH (1 N). (Kalweit et al., 2015) The final concentrations of Aβ were 100 μg/mL. The standard medium content (mg/L) were:Ca^2+^ 31.05; Mg^2+^ 17.6; Na^+^ 0.9; K^+^ 0.25; Fe^2+^ 0.001; HCO3^−^ 153.097; SO4^−^ 3; Cl^−^ 0.8; F^−^ 0.02; H_2_SiO_3_ 3.3 (pH=7.5). For *in vivo* investigations we applied Concanamycin A (ConA; 27689, 50 nM), 4,4′-dianilino-1,1′-binaphthyl-5,5′-disulfonic acid dipotassium salt (BisANS, D4162; 50 μM), propidiume-iodide (PI, 81845; 5 μM;), AO (13000; 15 μM) and NR (N-4638; 50 mM) dyes obtained from Sigma-Aldrich, USA. The rotifers were cultured based on the following methods of our previous publications (Olah et al., 2017). For standard food of cultures, we used homogenized baker’s yeast (EU-standard granulated instant form, 2-01-420674/001-Z12180/HU) which was heat-inactivated and filtered (Whatman filter with 10 μm pore; 6728-5100).

### Treatment and monitoring

Monitoring the size of the germovitellaria (Figure 1), in the presence of glucose (1 mM), started on Day 0 (D0) providing a reference control (Figure 2A and 2D). After twenty days (D20) of starvation (Figure 2B and 2E) and five days (D25) later the organ regeneration was presented (Figure 2C and 2F). On D20, one-time feeding (600 μg/mL yeast homogenate) was applied followed-by five days of recovery. The treatment protocol was performed on a ‘one-housed rotifer’ (one animal/well) setup in a 96-well plate (Costar Corning Inc., CLS3595). In all fluorescence-related experiments there were 150 ± 30 rotifer/well.

**Figure 1.**
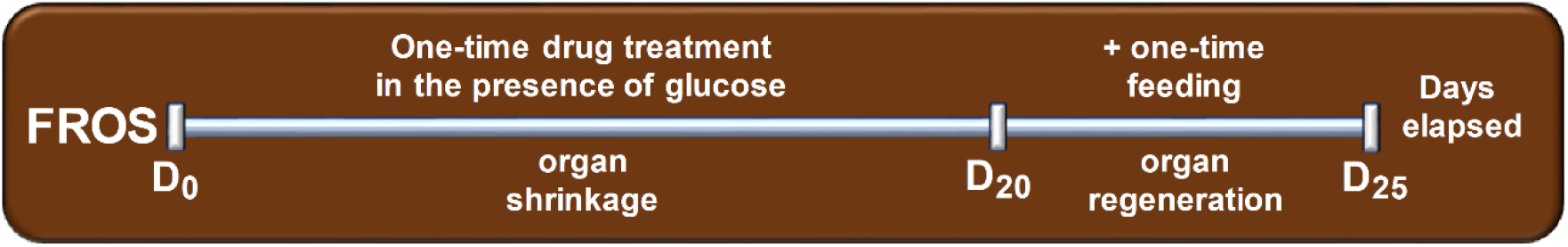
Schematic protocol of the functionally reversible organ shrinkage in a micro-invertebrate system. (D0 = Day 0, starting day; D20 = Day 20, organ shrinkage period with one-time drug treatment in glucose supplemented environment; D25 = Day 25, organ regeneration period with one-time standard feeding).

**Figure 2.**
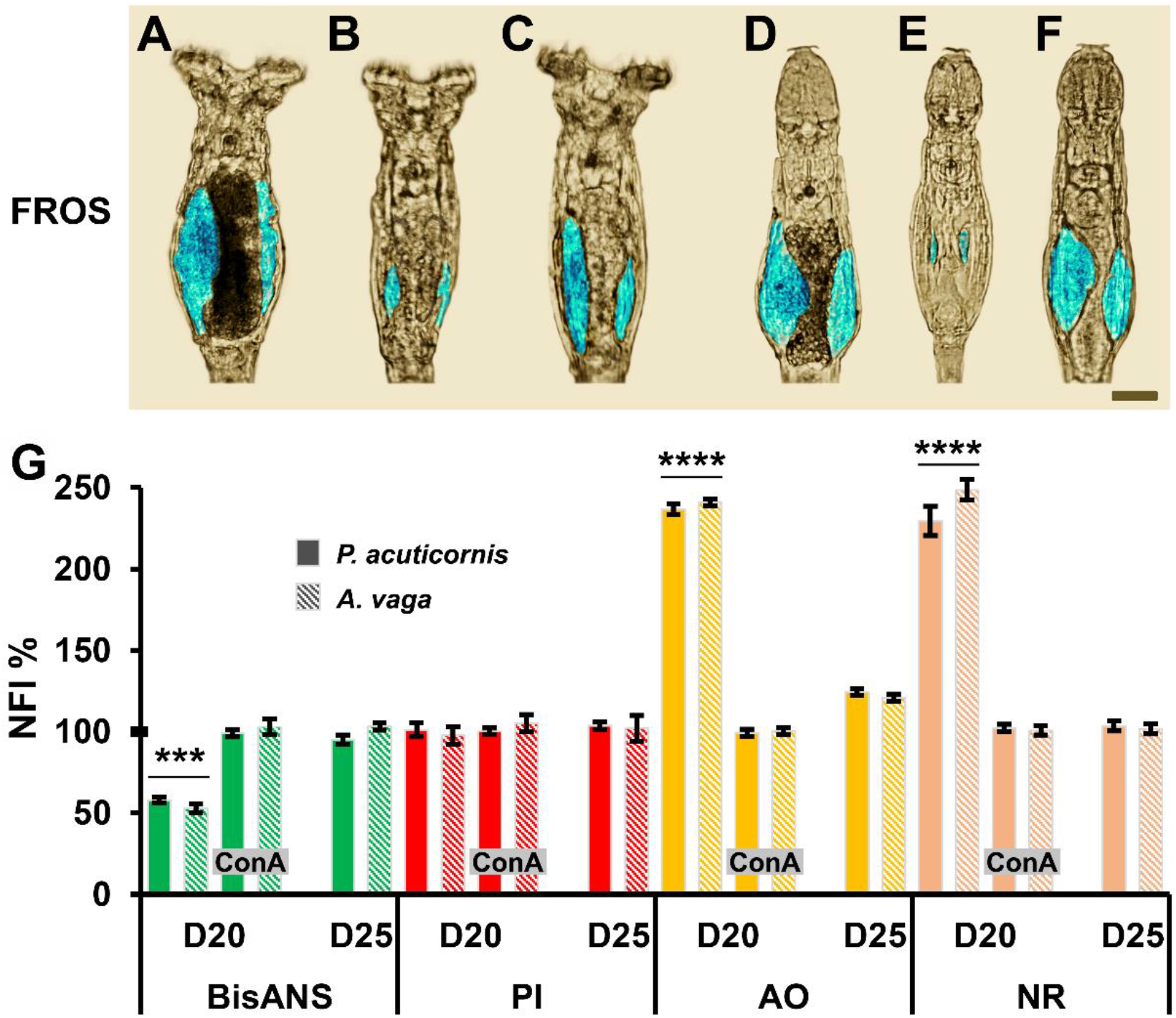
Functionally reversible organ shrinkage (FROS) in *P. acuticornis* and *A. vaga* rotifers. The germovitellaria (digitally pseudo-colored in blue; scale bar: 20 μm) of *P. acuticornis* (PA; A-C) and *A. vaga* (AV; D-F) was monitored on D0 (A and D), D20 (B and E) and D25 (C and F). The FROS-related alterations are presented (G) with the protein- (green), nucleic acid- (red) and AVOs amount in PA (full columns) and AV (striped columns) rotifers. The D0 means 100%. ConA was applied parallelly with all investigations. The error bars represent SEM. One-way ANOVA with Bonferroni *post hoc* test was used for statistical analysis, the levels of significance are p^***^ ≤ 0.001 and p^****^ ≤ 0.0001 (*, significant difference from all the groups).

The entities were photographed (Nikon D5100 camera, Japan) under the inverted microscope (Leitz Labovert FS). The two separated germovitellarium were digitally colored in blue (Figure 2A-F). The process of shrinkage with functional (egg production) regeneration capability was named ‘functionally reversible organ shrinkage’ (FROS).

To investigate (n=5, well) the amount of protein in the animals (Figure 2G), BisANS was applied (Datki et al., 2019) parallelly with detecting the total amount of nucleic acid, where PI was used (Mozes et al., 2011) after 10 min incubation and washing. The extinction/emission were 405/520 nm for BisANS and 530/620 nm for PI, measured by a microplate reader (NOVOstar, BMG, Germany).

In AVOs-detection methods, we used a slightly modified version of Kang et al. (2018). The AO and the NR labellings were performed under the same conditions as the BisANS and PI applications. The extinction/emission of AO was 480/620 nm in red and 480/520 nm in green. The wavelengths of NR were 540/630 nm in red and 450/590 nm in yellow. In order to inhibit the autolysis-related processes, ConA was added to the wells on D0 (Figure 2G). The percentage of normalized fluorescence intensity (NFI%) was calculated from the ratio of fluorescence data divided by the number of rotifers in each well.

In the experiential monitoring of FROS (Figure 3A), each data (n = 36; individual one-housed rotifer/well) sums the whole (bilateral) size of the germovitellaria in one individual. The organs were circled by using a freehand tool (allowing to create irregularly shaped selections) in ImageJ program (64-bit for Windows, (Collins, 2007). The scale bar was 20 μm.

**Figure 3.**
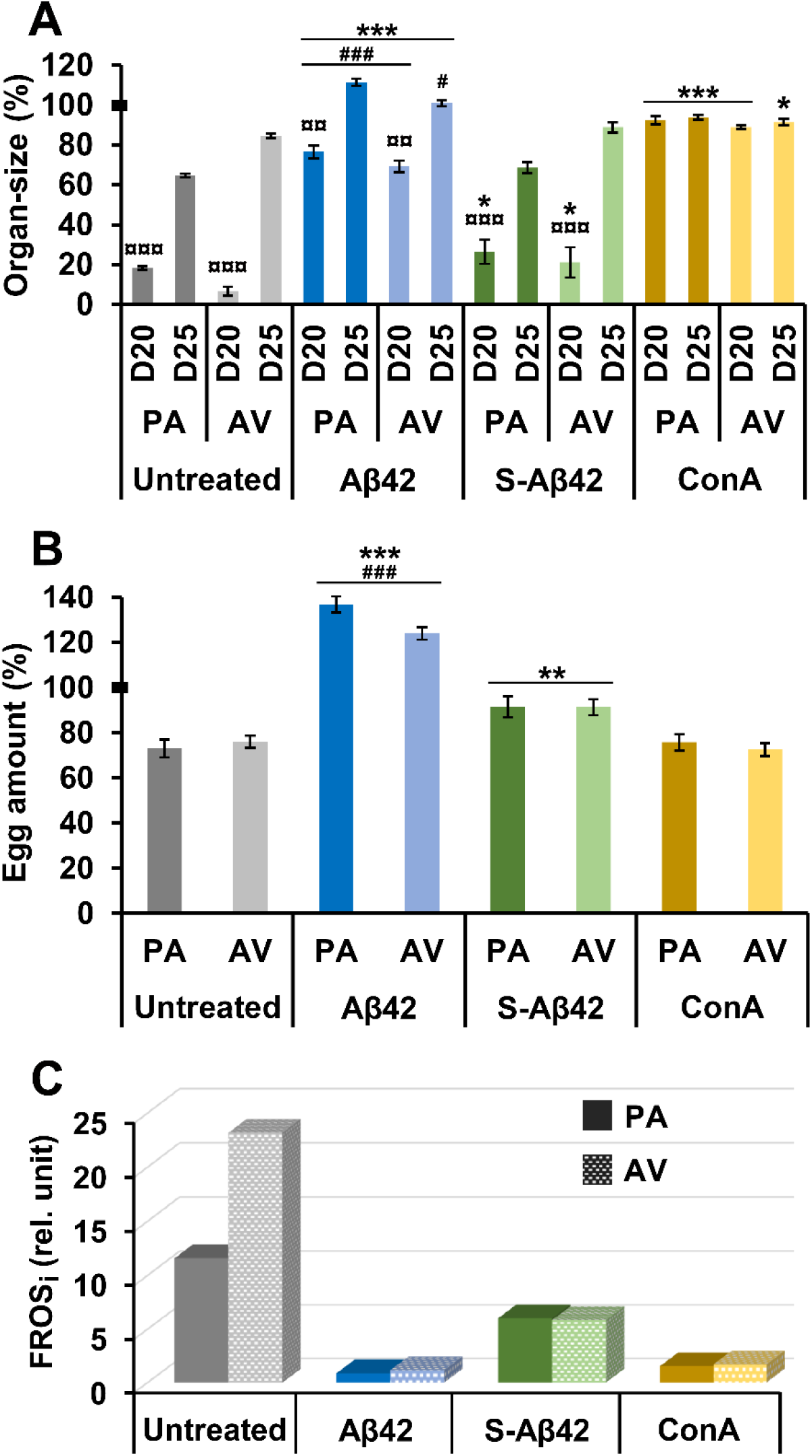
Modulation of the functionally reversible organ shrinkage (FROS) by aggregated Aβ42. The FROS was assessed in both *P. acuticornis* (PA) and *A. vaga* (AV) with applying Aβ42, S-Aβ42 or ConA. The germovitellaria were monitored on D0, D20 and D25. (A) The organ-size during FROS (A) and the amount of eggs on D25 (B) are presented. The D0 means 100%. The error bars represent SEM. One-way ANOVA with Bonferroni *post hoc* test was used for statistical analysis, the levels of significance are p^*, #^ ≤ 0.05, p^¤¤^ ≤ 0.01 and p^***, ###, ¤¤¤^≤ 0.001 (*, significant difference from the same untreated control groups of the given rotifer species; #, significant difference from the same S-Aβ42-treated groups of the given rotifer species; **¤**, significant difference from the D0 and D25 groups of the given rotifer species). FROS index (FROS_i_; relative unit) is presented (C) in untreated control, Aβ42, S-Aβ42 and ConA groups.

To demonstrate the functional recovery of the reproductive organs (Figure 3B), the number of laid eggs was counted after the regeneration phase (from D20 to D25). As a reference control, the treatment-free species-specific laid egg production was determined after 5 days in normal culturing conditions.

### Statistics

Statistical analysis was performed with SPSS 23.0 (SPSS Inc, USA) using one-way ANOVA with Bonferroni *post hoc* test. The FROS index (FROS_i_) was calculated by the following formula: FROS_i_ = A/B*A/C*D/E (germovitellaria size on: A, D0; B, D20; C, D25; D, species-specific number of laid eggs under standard feeding; E, number of laid eggs on D25). The error bars represent the standard error of the mean (SEM). The different levels of significance are indicated as follows: p^*, #^ ≤ 0.05, p^**, ¤¤^ ≤ 0.01, p^***, ###, ¤¤¤^ ≤ 0.001 and p^****^≤0.0001 (*, significant difference from the untreated controls; #, significant difference from the same S-Aβ42-treated groups of the given rotifer species; **¤**, significant difference from the D0 and D25 groups of the given rotifer species).

## Results and discussion

The Aβ42 is a well-known neurotoxin, which is much prone to form highly-resistant aggregates in an aquatic environment (Lin et al., 2019). The bdelloid rotifers are able to catabolize these aggregates, with no physiological damage (Datki et al., 2018). Our aims were to reveal the special role of Aβ42 in autocatabolism-related processes during a 25-day period (Figure 1). To explore the possible sequence specificity of this molecule, we applied their scrambled version as control. The molecular content and weight of S-Aβ42 were the same as that in wild-type human form, with different order of amino acids (Datki et al., 2018; Bartus et al., 2018).

On D0, the reference germovitellaria of *P. acuticornis* (Figure 2A) or *A. vaga* (Figure 2D) can be seen. Their size and fine structure reduced after a 20-day long glucose-supplemented (ATP-source for autolytic processes) starvation (Figure 2B and 2E). On D20, the animals were fed once, providing standard nutrient for the regeneration phase. The organs were then rebuilt and showed similar characteristics (Figure 2C and 2F) to the reference ones. These results suggest that the organic shrinkage is reversible depending on the availability of food. To connect the autocatabolism-related processes with starvation-induced organ shrinkage (Puente et al., 2016), we applied ConA (Johnson et al., 2010) during the experiments. The amounts of protein and nucleic acid were measured parallelly with AVOs. We found that the total protein decreased on D20, indicating that the animals catabolized it for survival. The ConA-administration inhibited these processes. The amounts of nucleic acid did not show any changes in either species (Figure 2G). In line with the above mentioned investigations we detected significant increase in autocatabolism-related vesicular acidification (Tan et al., 2018) in both species on D20 compared to the untreated reference values of D0. The ConA treatment hindered the observed alterations. On D25, there were no significant changes either in AO or in NR. The decrease of protein amount (associated with stable nucleic acid content) and the occurrence of AVOs under starvation show good conceptual correlation with organ shrinkage. These phenomena are adequate physiologic and experiential markers of autolytic metabolism in the aforementioned bdelloid rotifers.

The Aβs are stigmatized as negative multifunctional agents (e.g. in the human brain) in academic literature. Based on this conception, the core question is how Aβ42 may influence the autocatabolism-related processes in our microinvertebrate species? In FROS-related investigations the Aβ42, S-Aβ42 or ConA were added to the treatment medium on D0 (Figure 3A); therefore, these data served as references to the upcoming ones. In both rotifer species the organ shrinkage was less pronounced on D20 compared to the given untreated controls. Significantly higher regeneration was detected on D25 in Aβ42-treated groups compared to the other ones on D25. These results inferred that the native aggregate potentially has a specific modulatory effect. Decrease of organ-size was lower in Aβ42-treated groups compared to the ones influenced by S-Aβ42. The same phenomenon was observed at the end of regeneration, where the animals showed significantly higher rate of recovery in the presence of Aβ42. These data showed that the attenuation of shrinkage via modulation is likely sequence-specific, since the order of amino acid is the only difference between the two types of Aβs (Vadukul et al., 2017). On D20, ConA significantly inhibited the FROS in both species in a glucose-containing, but food-free environment. We have no knowledge about the treated animals eating the ConA itself, but the individuals remained alive in a good shape. On D25, the ConA had no effects on the monitored phenomenon.

In the FROS acronym, besides reversibility (‘R’) of organ shrinkage, ‘F’ stands for functionality, which refers to the remaining reproduction ability of rotifers. The measured parameter on D25 was the number of laid eggs, which was compared to the reference values (Figure 3B). In the presence of Aβs, the number of eggs significantly elevated in both species compared to the control and ConA peers; moreover, the Aβ42-modulated animals laid significantly higher number of eggs than the S-Aβ42-treated ones. Integration of egg-number-data into the time-dependent organ size alterations resulted a special formula, named FROS index, which is a representative unit for the current treatment agents. This index shows a linear correlation with the level of autocatabolism. The FROS_i_ was positively lower in all agent-influenced experiments compared to the controls in both species (Figure 3C). The FROS_i_ of native aggregates, in contrast to their scrambled version, demonstrated that the Aβ42 is not only a food source for rotifers, but it is also a potential regulator of their systemic metabolism. Both *Philodina* and *Adineta* species showed alterations with the same tendencies; therefore, these effects of Aβ42 are not limited to only one species.

The starvation-induced shrinkage of germovitellaria, with their regeneration and reproduction capability in these animals, is an adequate physiologic and experiential marker of autocatabolism, summarized by FROS_i_. By applying these microinvertebrates, the hitherto unknown roles of Aβ42 were demonstrated, providing additional tools for exploring relations between neurotoxic aggregates and metabolism.

## Acknowledgements

The authors wish to thank to Anna Szentgyorgyi MA, a professional in English Foreign Language Teaching for proofreading the manuscript.

## Author contribution

Conceptualization, Z.G.O. and Zs.D.; Methodology, Zs.D. and Z.G.O.; Validation, Zs.D., Z.G.O., and E.B.; Formal analysis, Zs.D., Z.G.O. and B.G.; Investigation, Zs.D., Z.G.O., B.E., B.G. and Zs.B. Resources, Zs.D., B.G. and J.K.; Data curation, Zs.D. and Z.G.O.; Writing—original draft preparation, Zs.D. and Z.G.O.; Writing—review and editing, Zs.D., Z.G.O., E.B., B.G., Zs.B. and J.K.; Visualization, Zs.D.; Supervision, Zs.D. and Z.G.O.; Project administration, Zs.D., Z.G.O., E.B., and Zs.B; Funding acquisition, B.G. and J.K.

